# Percolation and lifestyle transition in microbial metabolism

**DOI:** 10.1101/2025.01.12.632617

**Authors:** Rydberg Supo-Escalante, Germán Plata, Dinara R. Usmanova, Dennis Vitkup

## Abstract

Microbial species exhibit a remarkable diversity in their metabolic properties, genome composition, and ecological distribution. A central challenge of systems biology is to understand the relationships between genomic, metabolic, and phenotypic properties of bacteria. However, it is currently not well understood how the structure of metabolic network defines and reflects the lifestyle of diverse bacterial species across the tree of life. By analyzing thousands of genome-scale metabolic models of bacteria, we found a percolation-like transition in their ability to grow on independent carbon sources at around 800 metabolic reactions or about 2000 protein-coding genes. The observed transition is characterized by significant changes in metabolic network functional connectivity primarily associated with the completion of central carbon metabolism and the TCA cycle. Strikingly, experimentally observed phenotypic properties of bacteria also exhibit two markedly different regimes below and above the transition. Species with metabolic network sizes below the transition are typically obligate symbionts and require complex minimal media for their growth. In contrast, species with networks above the transition are primarily free-living generalists. The observed percolation transition is also reflected in multiple other genomic properties, such as a substantial decrease in the fraction of regulatory genes below the transition and higher evolvability for new metabolic phenotypes above the transition. Furthermore, we find that the distribution of bacterial genome sizes from unbiased environmental metagenomic sequencing also reflects genomic clusters corresponding to the observed transition. Overall, our work identifies two qualitatively different regimes in microbial metabolism and lifestyles characterized by distinct structural and functional properties of their metabolic networks.

## Introduction

The prokaryotic genome sizes vary considerably, spanning the range between a couple hundred kilobases (McCutcheon and Moran, 2012; Nakabachi et al., 2006) to well over ten megabases (Koonin and Wolf, 2008; Land et al., 2015). Bacteria are also remarkably diverse in their metabolic capabilities, including different energy acquisition strategies, and the ability to synthesize biomass through the assimilation of carbon and other elements from multiple organic and inorganic compounds (Berg, 2011; Madigan et al., 2012). Genomic and physiological diversity of bacteria is reflected in the structural and functional properties of their metabolic networks which allows microbes to live across many different environments and microbial communities. Comparative genomics and analysis of genome-scale metabolic models (GEMs) were previously used to study metabolic networks properties and evolution of bacterial phenotypes across the tree of life (Bentkowski et al., 2015; Nerima et al., 2010; Parter et al., 2007; Passi et al., 2021; Plata et al., 2015). Metabolic networks were also characterized by a scale-free power-law distribution of vertex connectivity (Barabási and Albert, 1999), by robustness to the failure of individual components across the three domains of life (Albert et al., 2000; Jeong et al., 2000; Smart et al., 2008), and by a modular and hierarchical structure (Csete and Doyle, 2004; Ravasz et al., 2002; Serrano et al., 2012). In prokaryotes, it has been demonstrated that obligate associations with a host often lead to massive genome reductions and to the loss of multiple biosynthetic pathways (McCutcheon and Moran, 2012; Murray et al., 2021). However, the exact structural changes and properties of metabolic networks responsible for different microbial lifestyles remain unclear.

An important property influencing the functionality of many real-world networks is percolation (Li et al., 2021). In physics, percolation refers to a geometric phase transition in a network, where the continuous addition of nodes or edges results in the sudden formation of a large connected network component (Yang, 2013). A percolation transition is characterized by a threshold, before which the network contains many connected clusters of small sizes, and after which a dominant giant connected cluster, comparable to the system size, starts to form. For metabolic network, biologically meaningful connectivity between chemical compounds can be defined by the feasibility of the flux between them, which can be determined using constraint-based methods like flux balance analysis (FBA) (Lewis et al., 2012; Orth et al., 2010). However, it remains unexplored whether percolation transition occurs in metabolic networks across different extant bacteria as their genome and network sizes grow. Furthermore, it is unclear what structural and functional properties of these networks may drive such percolation transition, and how the transition may influence phenotypes and lifestyles across diverse microbial species.

In this study, we investigated the role of metabolic network connectivity in shaping microbial metabolic phenotypes. First, we constructed thousands of GEMs across all bacterial genera. Using FBA, we examined the carbon sources that can support growth and biomass synthesis and then explored the global connectivity properties of metabolic networks. We correlated the ability of metabolic networks to interconvert various nutrients with available experimental data on culture media and symbiotic lifestyles. We identified the biochemical pathways whose completion usually drives a structural transition in the metabolic network connectivity. Finally, we explored the impact of this transition on how the number of genes in different functional categories scales with genome size and the distribution of genome sizes in environmental metagenomic samples. Overall, our study uncovered a percolation-like transition in metabolic networks across thousands of bacterial species, linking this phenomenon to changes in multiple functional and genomic properties, including environmental versatility and lifestyle.

## Result

### Bacterial growth phenotypes and metabolic network size

For bacterial species, the ability to metabolize different nutrients is a basic but crucially important phenotype which determines their fitness, lifestyles, and environmental distribution (Pudlo et al., 2022; Zeng et al., 2022). To investigate the relationship between the ability of prokaryotes to utilize different nutrients for growth and their metabolic network size, we considered 3,068 metabolic models of bacterial species reconstructed using the ModelSEED pipeline based on genomes in the KBase platform (Arkin et al., 2018); to uniformly cover the bacterial phylogeny, in our analysis we considered a single or a few representative reconstructions per each bacterial genus (see **Methods**). We then used FBA to predict, for each metabolic model, how many potential carbon sources can individually support biomass synthesis when coupled with a trace of other carbon compounds and unconstrained flux of non-carbon compounds (see **Methods**). Notably, the comparison across bacteria between the number of different carbon compounds that can sustain growth and the corresponding metabolic network sizes, i.e., the number of reactions in each metabolic network, showed two different regimes (**Fig. 1a**). Specifically, for metabolic networks with more than ∼800 reactions, the number of carbon compounds that can support biomass synthesis increases approximately linearly with the number of network reactions, while below this threshold, very few carbon compounds provided individually were predicted to support biomass synthesis in the corresponding microbial species.

**Figure 1.**
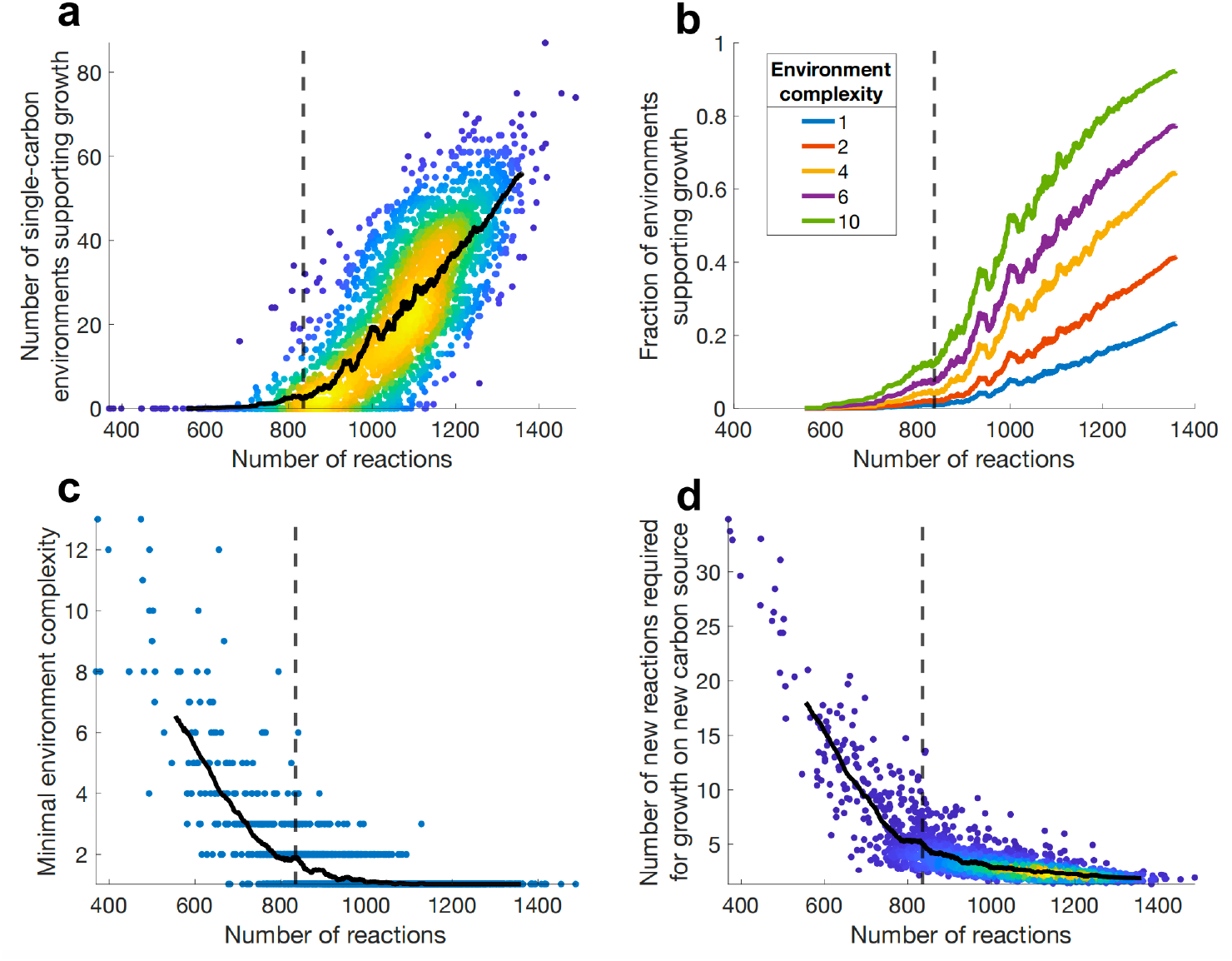
Growth on single and multiple carbon sources and phenotypic evolvability of bacterial metabolic networks. Each dot represents a metabolic model for different bacterial species (N = 3,068); metabolic network sizes are measured as the number of reactions in the corresponding models. Black lines show a 100-reaction moving average of the data. Dashed lines represent the fitted percolation threshold **(Methods). a**, Number of different single-compound carbon sources (out of 227 tested) that can support biomass synthesis. **b**, The fraction of metabolic environments of different complexity that can support biomass synthesis; the environmental complexity represents the number of carbon compounds provided simultaneously. **c**, Minimum carbon environment complexity needed for biomass synthesis as a function of metabolic network size. Minimum complexity represents the smallest number of carbon compounds needed to be provided simultaneously to support biomass synthesis. **d**, Average number of additional reactions required to enable biomass synthesis on a new single-compound carbon source (for 30 randomly tested carbon compounds) as a function of metabolic network size.

To further explore the relationship between metabolic network size and biomass synthesis, we investigated the bacterial growth in environments where multiple carbon sources are present simultaneously (see **Methods**). To that end, we defined the complexity of each environment as the number of individual carbon compounds that are simultaneously present in the environment. Using FBA, we then calculated the ability of microbial metabolic networks to synthesize biomass in environments of different complexities **(Fig. 1b)**. As expected, environments with higher complexity, i.e., more available carbon sources, generally enabled a larger fraction of metabolic models to synthesize biomass. Interestingly, similar to growth on single carbon sources **(Fig. 1a)**, the number of carbon environments enabling biomass synthesis significantly increased for bacteria with over ∼800 reactions, across different environmental complexities **(Fig. 1b)**. We also explored the minimum environmental complexity required for metabolic networks of different sizes to enable biomass synthesis **(Methods)**. Notably, this analysis identified two regimes, again with a transition at ∼800 reactions **(Fig. 1c)**, with networks larger than this threshold typically being able to synthesize biomass on a single or few carbon compounds, while networks below the threshold usually required multiple carbon sources to be simultaneously provided for growth.

Evolvability, defined as the ability to acquire new phenotypes, for example through horizontal gene transfer of enzyme-coding genes, is crucial for bacterial survival when facing diverse environmental conditions (Power et al., 2021). Thus, we investigated how the evolvability of metabolic bacterial phenotypes depends on metabolic network size. To that end, for each metabolic model we used Mixed Integer Linear Programming (MILP) to determine the minimum number of additional metabolic reactions required, on average, to enable biomass synthesis on a new single-compound carbon source **(Methods)**. Interestingly, while metabolic networks with more than ∼800 reactions could synthesize biomass on new single-compound carbon sources with 5 or fewer additional reactions, this number sharply increases to 10 or higher for networks with less than ∼800 reactions (**Fig. 1d**). This result suggests that metabolic networks with less than ∼800 reactions are also significantly less evolvable towards new metabolic phenotypes compared to network above that threshold.

### Formation of a giant connected component and metabolic network percolation

The results presented above suggests that living bacteria with network sizes below ∼800 reactions usually can synthesize biomass only in a relatively small number of different environments and that they often require for growth several carbon compounds to be provided at the same time. These bacteria are also significantly less evolvable because they need a substantially higher number of additional reactions to be able to utilize a new carbon source for biomass synthesis. Given these intriguing results, we next asked why the relationship between the bacterial growth phenotypes and metabolic network size exhibits the two distinct functional regimes, i.e. above and below ∼800 reactions, as well as the nature of the transition between these regimes. Because biomass synthesis crucially depends on the flux of mostly carbon-containing compounds through the metabolic network, from nutrients to biomass precursors, we next explored the ability of bacteria to interconvert different metabolites into each other. Specifically, we defined a directed edge between two carbon-containing compounds when one of them could be, as predicted by FBA (**Methods**), synthesized from the other. We then built a metabolite interconversion graph, with edges connecting metabolites that can be mutually interconverted, and calculated for each considered bacteria the largest set of metabolites that can all be synthesized from each other. Similar to the results describing the number of independent carbon sources supporting biomass synthesis (**Fig. 1a**), we found that the size of the largest connected network component increases substantially faster for bacterial metabolic networks larger than ∼800 reactions **(Fig 2a)**. Such a sharp increase in the network connectivity is typically associated with a phenomenon known in the network theory as a percolation transition (Li et al., 2021). Based on a linear-power law fitting, we determined that the percolation critical point corresponds to a metabolic network size of 820 ± 70 reactions (see **Methods**). Naturally, the fast increase of the largest connected component above the transition should facilitate bacterial growth on multiple metabolites as the network connectivity promotes the interconversion of various nutrients into common biomass precursors (**Fig. 1a**).

**Figure 2.**
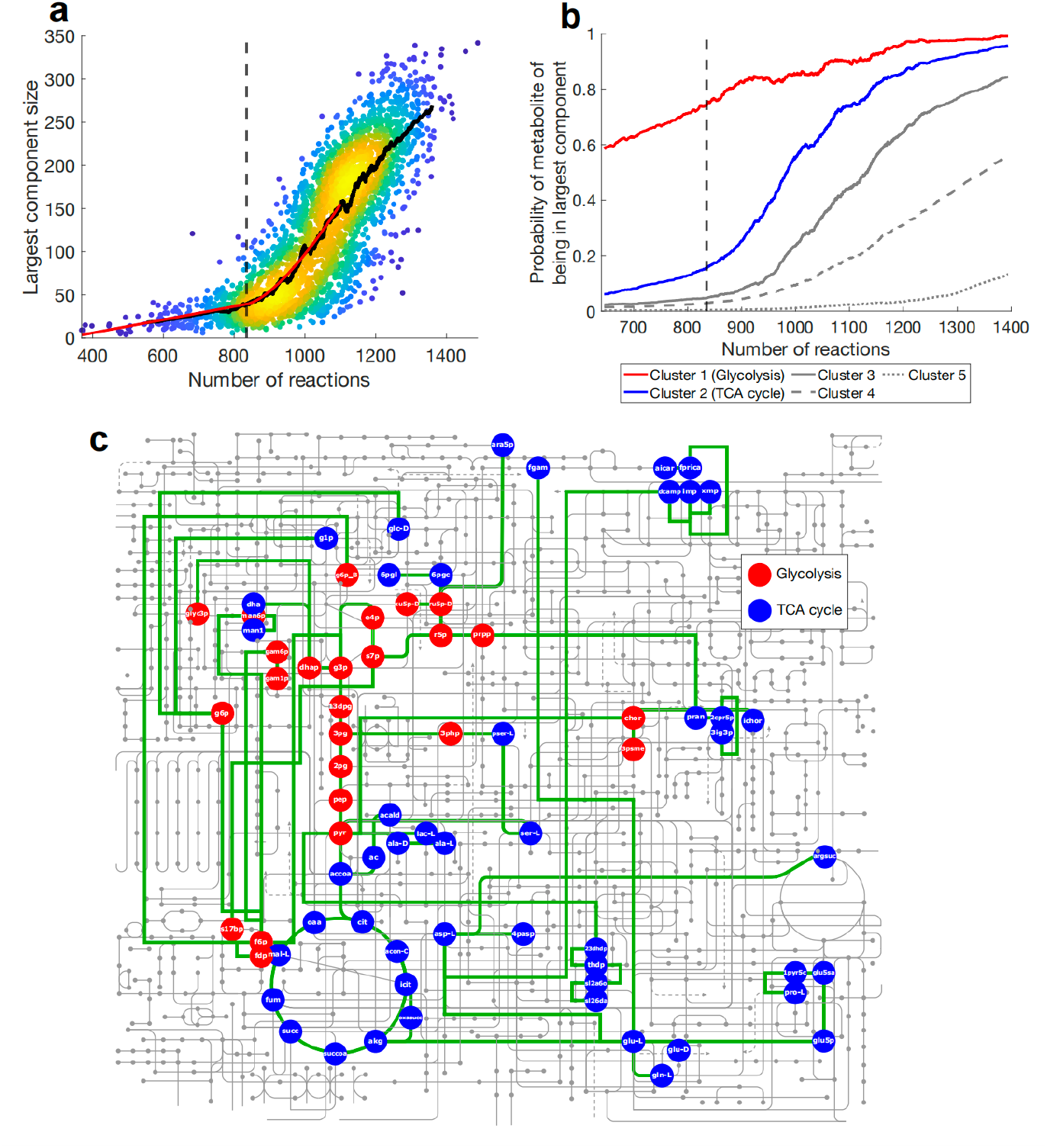
Percolation transition in metabolic network connectivity and the nature of the largest component. **a**, The size of the largest component in bacterial metabolite interconversion graphs as a function of the corresponding metabolic network size. The dashed black line shows the fitted percolation transition threshold (820 ± 70). The black line shows the average across a 100-reaction sliding window over metabolic network sizes. The red line represents a linear-power law fit used to determine the approximate percolation threshold **(Methods). b**, Probability profiles of carbon compounds to be a part of the largest connected network component as a function of metabolic network size. Profiles were clustered into five different groups using pathway analysis (**Methods**). **c**, Metabolic map highlighting metabolites related to the glycolysis (red), and the TCA cycle (blue) that correspond to the clusters of the same color in **b**.

We next investigated the biochemical pathways whose completion commonly coincides with the observed percolation in the connectivity of the metabolic network. To that end, we calculated the probability of each carbon compound to be a member of the largest connected component for bacteria with different network sizes **(Methods). (Suppl Fig 1)**. We then performed K-means clustering to group metabolites with similar probability profiles across metabolic networks sizes and used pathway analysis from the MetaboAnalyst 6.0 web-based platform (Pang et al., 2024) to identify biochemical pathways primarily associated with each cluster **(Fig. 2b)**. The first cluster (red color in **Fig. 2b**) contains metabolites that have, even below the percolation transition, the highest probability of being in the largest component. All metabolites in this cluster belong to glycolysis and the pentose phosphate pathway (PPP) **(Fig. 2c, red)**. Notably, metabolites from the second cluster demonstrate the sharpest increase in the probability of being in the largest network component around the percolation threshold, and have very high probabilities to be in the largest connected component above the transition. They mostly participate in the tricarboxylic acid (TCA) cycle and associated reactions **(Fig. 2c, blue)**. Metabolites from other clusters **(Fig. 2b, gray)** demonstrate a relatively slower increase in their probability to be in the largest component; the metabolites in these clusters are primarily associated with secondary metabolism **(Supplementary table 1)**. These results demonstrate that bacteria around the percolation transition share a universal core of metabolites in the largest component of their metabolic interconversion graph. This core generally includes metabolites from the glycolysis, the PPP and the TCA pathways.

### The influence of the metabolic percolation transition on lifestyle and genome composition in bacteria

Our analyses of structural and functional properties of metabolic networks demonstrate qualitatively different behavior below and above the percolation-like transition. To complement these results, we next asked whether the observed changes in the network properties correlate with significant alterations in bacterial lifestyles and nutritional requirements observed experimentally. To explore this, we first utilized the KOMODO database (Oberhardt et al., 2015), which describes media ingredients supporting the growth of thousands of microbial strains; the KOMODO database is specifically focused on complex media ingredients that comprise a rich mixture of compounds (e.g., yeast, plant or meat extracts). Interestingly, we observed a significant increase in the average number of complex media ingredients required to culture species with metabolic networks containing fewer than ∼800 metabolic reactions (**Fig. 3a**). These results are consistent with our computational results (**Fig. 1, 2**) and demonstrate that substantially more complex media is usually required for growth of bacteria with metabolic networks below the transition (Zarecki et al., 2014). Next, we also analyzed the distribution of obligate symbiotes across bacteria with different network sizes (**Fig. 3b**). To that end, we used data describing lifestyles for ∼700 bacterial species (Burstein et al., 2016). We found that the prevalence of obligate symbiotes does not increase gradually with decreasing metabolic network size. Instead, there is a sharp increase in the frequency of symbionts near the network percolation threshold of about ∼800 reactions (**Fig. 3b**). Overall, these experimental results demonstrate that significant changes in the bacterial metabolic network structural properties coincide with qualitative differences in the physiological and lifestyle properties of bacteria observed below and above the percolation transition.

**Figure 3.**
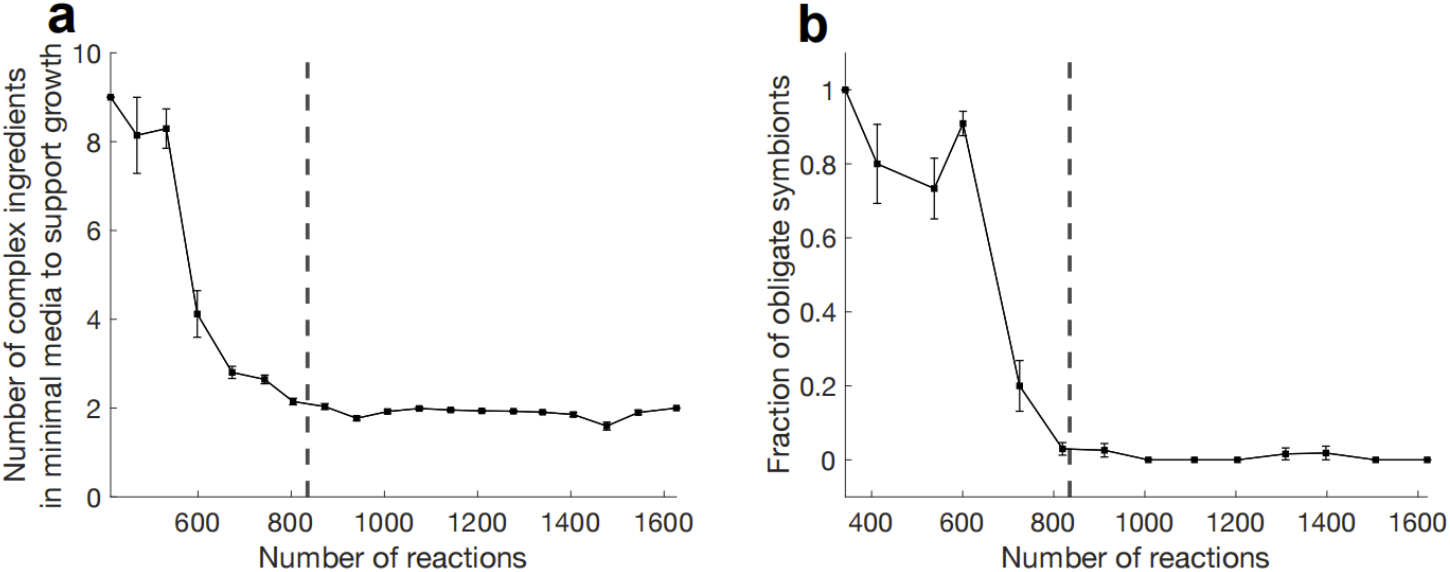
Nutritional and lifestyle correlators of the percolation transition in metabolic network connectivity. **a**, The number of complex media ingredients in the simplest known culture media for different bacteria as a function of metabolic network size. Data on media composition were obtained from the KOMODO database (Oberhardt et al., 2015), complex media ingredients, as defined in Oberhardt *et al*. (Oberhardt et al., 2015). **b**, The fraction of obligate symbionts among species with different metabolic network sizes. Lifestyle classifications were obtained from the study by Burstein *et al*. (Burstein et al., 2016). In both panels, error bars represent the SEM. Number of reactions were obtained from metabolic networks of the corresponding species using the KBase resource (Arkin et al., 2018). The dashed lines show the approximate percolation threshold.

The divergence in bacterial lifestyles and metabolic versatility at the percolation transition should also correlate with other functional properties of prokaryotic species. To investigate this, we explored the relationship between the number of genes associated with several distinct functional categories and the total number of genes in bacterial genomes. First, using genomes with annotation available from the Clusters of Orthologous Groups (COG) database (Galperin et al., 2021), we selected bacterial gene families representing housekeeping genes involved in universal processes such as transcription, translation, replication, RNA processing, and protein turnover, as well as families representing environmental genes, including signaling proteins, mobile elements, and defense genes. Second, for genomes used in the analyses of network connectivity, we calculated the number of transcription factors predicted using profile searches with hidden Markov Models (HMMs) in the PFAM database (Mistry et al., 2021) **(see Methods)**. Consistent with reported observations (Molina and van Nimwegen, 2009; Sela et al., 2019), genes in each of these functional and molecular categories showed a power-law scaling with the total number of genes in bacterial genomes **(Fig. 4)**. Interestingly, for bacterial genomes with fewer than ∼2000- 2300 coding genes, which approximately correspond to the ∼800 metabolic reaction percolation threshold **(Methods, Suppl Fig 2)**, the scaling for environmental genes and for transcription factors becomes substantially steeper when genome sizes decreased below the transition threshold **(Fig. 4b, c)**. In contrast, for housekeeping genes the scaling becomes slightly less steep when crossing the threshold **(Fig. 4a)**. For all functional categories the piecewise linear regression with two different power law scaling anchored at the percolation threshold **(see Methods)** explained the data better than a single power law across all genome sizes (power-law slope change > 0.5, F-test p-value < 10^−6^). Altogether, these results suggest that microbes with metabolic networks below the transition are usually associated with less complex gene regulation (Cottrell and Kirchman, 2016), a smaller fraction of genes associated with environmentally regulated processes (Sela et al., 2019), and a higher fraction of genes associated with housekeeping functions. These patterns are consistent with the hypothesis that bacteria with networks below the transition typically live in more stable and symbiotic environments.

**Figure 4.**
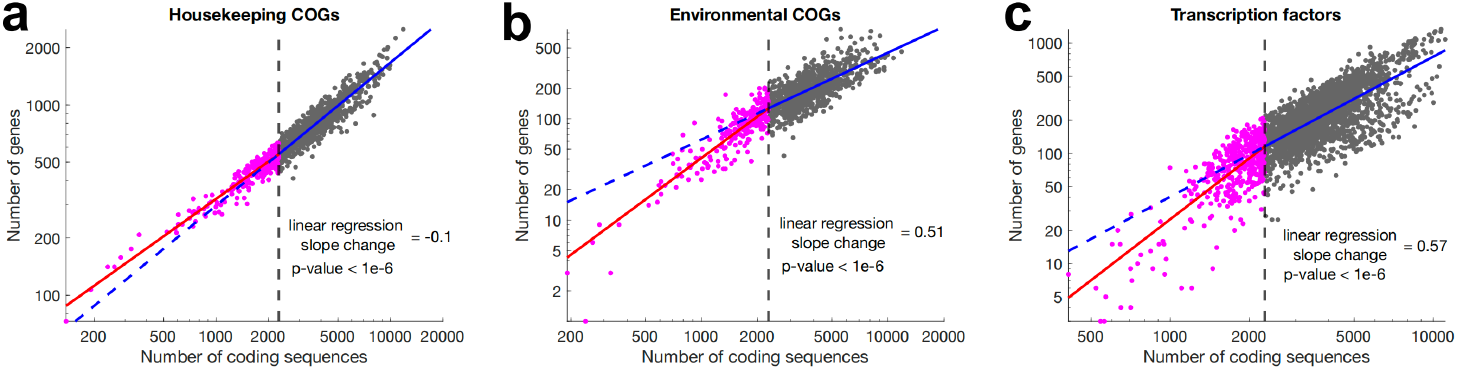
Scaling of the number of genes in specific functional and molecular categories with the total number of protein-coding bacterial genes. The number of genes annotated in three different functional and molecular categories as a function of the total number of protein-coding bacterial genes. **a**, Housekeeping, and **b**, Environmental genes from bacterial COGs (Galperin et al., 2021). **c**, Genes predicted to be transcription factors from bacterial genomes used in the connectivity analyses. The red and blue lines represent a two-segment piecewise linear fit of the log-transformed data (see **Methods**). The vertical dashed lines indicate the approximate percolation threshold (∼2300 protein-coding genes) (**Suppl Fig 2**). The slope change indicates the difference in slope above and below the percolation threshold.

Finally, we investigated whether the distribution of genome sizes of extant microbes also shows an imprint of two distinct regimes in their metabolic network and lifestyle properties. Because across prokaryotes, the number of coding genes closely correlated with the genome size, with one gene approximately corresponding to one genomic kilobase (Konstantinidis and Tiedje, 2004), the percolation transition approximately corresponds to a genome size above ∼2 Mb. To explore the distribution of genome sizes, we used a recently published metagenomic dataset with estimated genome sizes for species from terrestrial and aquatic samples (Rodríguez-Gijón et al., 2022). Metagenomic sequencing allows for the retrieval of genomes from all microorganisms present in an environmental sample (Hugenholtz and Tyson, 2008), and microbes with complex nutritional requirements, which are often difficult to culture in the lab, are expected to be well represented in metagenomics samples. Thus, metagenomic samples should provide an unbiased representation of the bacterial genome size distribution in a particular environment. The distribution of microbial genomics sizes from both aquatic and terrestrial samples suggested a bimodal trend. Therefore, we used a gamma mixtures model with two components to fit the genome size distributions. For aquatic **(Fig. 5a)** and terrestrial **(Fig. 5b)** samples, we found the existence of two peaks around ∼1.4 Mb and ∼3.4 Mb in the distribution of genomic sizes (1.4 Mb and 3.3 Mb for aquatic and 1.3 Mb and 3.8 Mb for terrestrial). Notably, the approximate percolation genome size threshold (2.3 Mb, indicated by vertical dashed lines in **Fig. 5**) is located in between the two modes of genome size distribution in both the aquatic and terrestrial metagenomic samples. Overall, these results suggest that microbial species from the unbiased environmental samples form genomic clusters with peaks on each side of the metabolic and lifestyle percolation transition.

**Figure 5.**
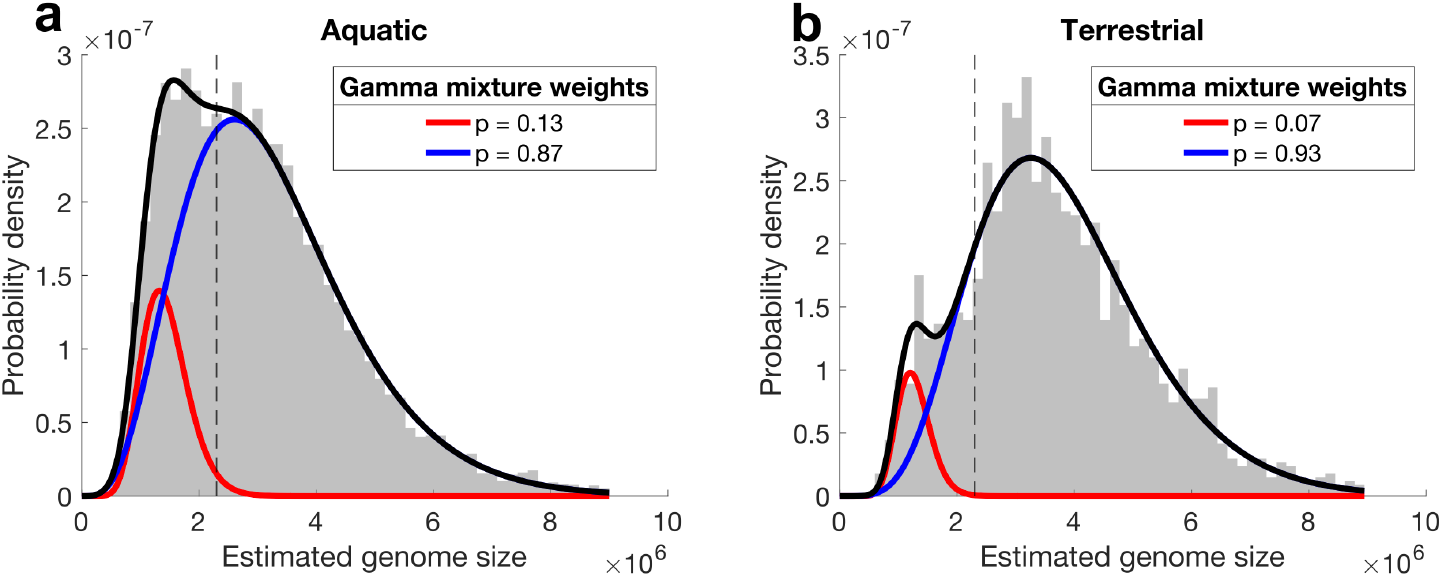
Distribution of microbial genome sizes based on metagenomic sequencing. The figure shows the distribution (grey) of estimated genomes sizes for metagenomic samples from **a**, aquatic, and **b**, terrestrial environments (Rodríguez-Gijón et al., 2022). In each case, a gamma mixture model with two components (red and blue) was fitted to the distribution data. The dashed vertical line indicates the genome size approximately corresponding to the percolation transition. The black line represents the total gamma mixture model fitted to the experimental data.

## Discussion

Our study demonstrates a fascinating connection between a previously unknown percolation-like transition in the global structural connectivity of bacterial metabolic networks and the corresponding microbial lifestyles and phenotypes. We found that functional and connectivity properties of bacterial metabolism do not scale smoothly with genome and network sizes. Instead, we identified two qualitatively different regimes across bacterial metabolic networks of various sizes. Metabolic networks with sizes below the transition remain relatively fragmented and specialized, while above the transition we observe a rapid growth in the overall network connectivity when disparate metabolic pathways usually merge into highly connected and flexible networks. Similar to observations in other real-world networks (Artime et al., 2024), crossing the transition usually leads to the formation of a large connected component, enabling bacteria to efficiently interconvert a variety of metabolites into each other. The ability to interconvert multiple metabolites and nutrients is likely to be particularly important for free-living bacteria inhabiting diverse metabolic environments, which may explain their predominance above the transition. Although it was previously observed that many host-dependent microbial symbionts have genomes smaller than 1 Mb (McCutcheon and Moran, 2012), the percolation-like transition we found occurs at approximately 2 Mb, indicating that free-living bacteria usually have substantially larger genome and network sizes.

It is important to emphasize that real bacterial metabolic networks are not random but are shaped by biological evolution, which endows them with unique structural and functional properties. Notably, an intrinsic characteristic of cellular metabolism, as previously described (Csete and Doyle, 2004), is its hourglass-like structure. This structure enables networks to interconvert diverse environmental nutrients into a common set of intermediate core metabolites, such as Acetyl-CoA or pyruvate, and then use these core metabolites to synthesize various biomass components. Supporting this conceptual framework, we found that the completion of the TCA cycle, along with several associated pathways, such as glycolysis and the pentose phosphate pathway, plays a major role in the observed percolation transition. This finding underscores that, beyond its central role in aerobic energy production, the TCA cycle also functions as a crucial biosynthetic hub, supplying essential precursors for the biosynthesis of numerous amino acids, lipids, and vitamins (Stobbe et al., 2012). Consistent with our results, bacterial symbionts are frequently observed to have incomplete TCA cycles (González-Domenech et al., 2012; Herlemann et al., 2009; Lin et al., 2024). Intriguingly, the TCA cycle likely emerged early in the evolution of life on Earth (Nunoura et al., 2018), as some of its reactions are feasible without enzymatic catalysts (Keller et al., 2017). After its emergence, the TCA cycle likely facilitated the rapid diversification and adaptation of microbial metabolism for growth in different environmental conditions.

Our analysis demonstrates that globally connected metabolic networks above the transition enable substantially higher phenotypic evolvability, in particular the ability to gain growth on a new carbon source. This increased evolvability likely arises from incorporating necessary new metabolic reactions into a highly connected network core (Maslov et al., 2009). Previously, we described the microbial phenotypic clock, i.e., the continuous diversification of bacterial metabolic phenotypes overs billions of years across the microbial tree of life (Plata et al., 2015). It is likely that globally connected metabolic networks facilitate this diversification and adaptation to new environments. In contrast, adaptation to more stable and symbiotic metabolic environments likely result in substantially less connected networks. Indeed, faster rates of gene loss, driven by both selection and drift (Fisher et al., 2017), has been observed during the evolution of obligate symbiotic bacteria (Burke and Moran, 2011; Toh et al., 2006; Wolf and Koonin, 2013). Rich environments have been also shown to promote the rapid evolution of multiple auxotrophies in experimental evolution assays (D’Souza and Kost, 2016). Moreover, adaptation to more stable and restricted environments below the transition is consistent with our observation of smaller fractions of transcriptional regulators and genes responsible for environmental interactions. We note that a substantial fraction of bacterial clades, such as the candidate phyla radiation (CRP) (Brown et al., 2015; Hug et al., 2016), are likely to have metabolic networks that lie predominantly below the transition. In the future, it would be interesting to further investigate how metabolic properties of bacteria above and below the transition influence their distinct functional traits, interaction strategies, and ecological niches.

## Methods

### Species selection and metabolic model reconstruction

Starting with bacterial genomes available on the KBase platform (Arkin et al., 2018), we retained a single representative genome from each bacterial genus. Each representative genome from a genus was selected based on having the median genome size within that genus. The selected genomes were annotated using RAST (v1.9.1), and genome-scale metabolic models (GEMs) were then reconstructed using the “Build Metabolic Model” method (v2.0.0) in the KBase Narrative interface (Arkin et al., 2018), which implements the ModelSEED pipeline (Henry et al., 2010). We excluded GEMs from the order *Enterobacterales* because their genomes had a substantially higher number of metabolic annotations and larger metabolic networks compared to other bacteria with similar genome sizes—a likely consequence of annotation transfers from the well-annotated *Escherichia coli* genome. This process resulted in a dataset of 3,068 bacterial GEMs used in the analyses.

Additionally, for the analysis of bacterial nutritional and lifestyle properties we built 5,067 and 672 GEMs, respectively, using the aforementioned procedure for species found in the KOMODO database (Oberhardt et al., 2015) and the database of lifestyle classifications (Burstein et al., 2016). For the analysis of genome functional categories, we built 1,310 GEMs for species listed in the COGs database (Galperin et al., 2021), using genomes with corresponding NCBI genome database identifiers.

### Model processing

For the subsequent analyses involving FBA, several modifications and adjustments were made to the reconstructed GEMs. First, all sink reactions were made irreversible to prevent the unrestricted intracellular uptake of metabolites. This ensured that nutrients were only taken up from the extracellular compartment. Second, demand reactions were added for every metabolite in the GEMs. Since demand reactions only remove metabolites from networks, they did not alter the metabolic network topology. However, demand reactions allowed the GEMs to consume intracellular metabolites in order to balance metabolites that accumulated in the networks. Third, reactions that were not properly mass-balanced in terms of carbon atoms were blocked, with two exceptions: (i) coupled reactions that are individually unbalanced for carbon but balanced as a pair, and (ii) reactions that are essential for the model to generate a non-zero biomass flux.

### Prediction of carbon sources supporting biomass synthesis

We compiled a list of 227 carbon-containing metabolites for which exchange reactions were present in at least one of the bacterial reconstructions. As described previously (Plata et al., 2015), for each model we determined whether each of the 227 compounds could support biomass synthesis when provided as the sole carbon source in excess. To achieve this, we used the COBRA toolbox (Heirendt et al., 2019) to perform the required FBA calculations. Specifically, for each metabolic reconstruction, we determined a baseline biomass flux using FBA (i.e., the maximum feasible flux through the biomass reaction) when the combined carbon uptake from all possible carbon compounds was constrained to less than 10 mmol g^−1^ dry weight per hour, while non-carbon compounds were provided in excess. Almost all models could achieve a baseline biomass flux of approximately 0.25 mmol g^−1^ dry weight per hour, characterizing the maximum biomass yield for the constrained model. Next, we calculated the maximum biomass flux when each tested compound was provided in excess (i.e., with a maximum uptake rate of 1 mol g^−1^ dry weight per hour), while the combined uptake of all other carbon compounds remained constrained to less than 10 mmol g^−1^ dry weight per hour. A carbon compound was considered capable of supporting biomass synthesis if the obtained biomass flux, when provided in excess, exceeded 0.5 mmol g^−1^ dry weight per hour—equivalent to double the maximum baseline biomass flux.

For environments of higher environmental complexity (i.e., when more than one carbon source was provided in excess simultaneously), a similar approach was used. In these scenarios, the uptake of each provided carbon source was limited to 1 mol g^-1^ dry weight per hour. The difference was that multiple carbon sources were supplied in excess at the same time. The total number of tested environments included either all possible combinations of carbon sources for the given complexity or 10,000 randomly sampled combinations. The additional complexities explored involved environments with 2, 4, 6, and 10 carbon sources provided in excess.

### Calculation of minimal environmental complexity required for biomass synthesis

To estimate the minimum number of carbon sources required to be provided simultaneously for biomass synthesis (defined as a biomass flux greater than 0.5 mmol g^−1^ dry weight per hour), we started by providing single carbon sources in excess and evaluating biomass synthesis with FBA. If at least one carbon source supported biomass synthesis, the minimum complexity was set to one. Otherwise, the environmental complexity was increased incrementally, and environments containing two different carbon sources in excess were tested for biomass synthesis using FBA, and the procedure was repeated until biomass synthesis was observed or all environments with complexity up to 15 were tested. For each environmental complexity a maximum of 10,000 random combinations of carbon sources were tested. The minimum complexity was then defined as the number of carbon sources in the first environment where biomass synthesis was observed. If no biomass synthesis occurred across environmental complexities, the minimum complexity was considered undefined.

### Prediction of reactions required to grow on a new carbon source

To estimate the smallest number of additional reactions required for growth on a new carbon source, we first created a global metabolic network by combining all reactions present in the individual GEMs. If a reaction was reversible in at least one of the individual reconstructions but irreversible in another, the reversible version was retained in the global reconstruction. Next, for each bacterial network, the analysis was performed as follows. First, for the corresponding reactions in the global reconstruction, the reversibility was adjusted to match the individual reconstruction. Second, the following constraints were applied to the global reconstruction: 1) the combined carbon uptake from the environment was set to less than 10 mmol of carbon g^−1^ dry weight per hour, and 2) the uptake of metabolites through exchange reactions that were not present in the individual reconstruction was set to 0. Third, we randomly selected 20 carbon sources that could not support biomass synthesis in the individual metabolic reconstruction. Fourth, for each of the selected possible new carbon sources, we formulated a set of distinct MILP problems in which each of the selected carbon sources were provided in excess, the biomass synthesis flux was constrained to be higher than 0.5 mmol g^−1^ dry weight per hour, and the optimization objective was set to minimize the number of additional metabolic reactions, i.e., reactions not present in the original individual reconstruction, required to achieve biomass growth.

### Metabolite interconversion graph

To identify connections between carbon metabolites in each network reconstruction, we began by blocking the uptake of all carbon compounds while keeping all other carbon-free compounds available in excess. For each pair of carbon compounds, we then tested whether they could be used to synthesize each other when provided as the only available carbon source. Specifically, for each compound in the pair, we added a source reaction and performed FBA to optimize the flux through the demand reaction of the other compound. If a feasible metabolic flux (greater than 10^−3^ mmol g^-1^ dry weight per hour) was observed in both directions, the pair of compounds was assumed to interconvert, and a connection was established between them.

To reduce computational time, interconversions were only calculated for metabolite pairs where one metabolite serves as a substrate and the other as a product in at least one reaction. By transitivity, any pair of metabolites not in the same reaction would still interconvert if a path of interconverted metabolites exists between them. Additionally, for metabolites containing the cofactors Coenzyme A (CoA) or Acyl Carrier Protein (ACP), following the previous approaches (Gao et al., 2021), the demand reactions were modified so that the cofactor portion of the compounds was recycled as a product. This adjustment was necessary because it is impossible for most GEMs to synthesize these cofactors.

### Largest connected component and approximate percolation threshold

For each metabolite interconversion graph of carbon compounds, we used the “conncomp” method in MATLAB to calculate all strongly connected components in the graph, i.e. all carbon metabolite subsets such that a directed path exists between all pairs of metabolites in the set, and no metabolite outside the set shares this characteristic. To determine the approximate percolation threshold, we fitted the size of the largest component as a function of the number of reactions using a linear-power law function (Stauffer, 1979):

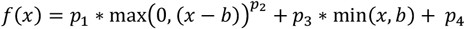

where, *x* represents the number of reactions, *b* is the percolation threshold, and *p*_1_, *p*_2_, *p*_3_, and *p*_4_ are the function parameters. This function was chosen because, immediately after the percolation threshold, the fraction of nodes in the largest component increases approximately following a power-law behavior. Accordingly, only bacterial GEMs with at most 1,100 reactions were considered in the fits, and the percolation threshold was searched within the range of 500 to 1,000 reactions.

To perform the fitting, we used the “lsqcurvefit” method in MATLAB with the starting parameters p_1_=0.1, p_2_=1.5, p_3_=0.07, and p_4_=-23. The lower and upper bounds for each parameter where set to −∞ and to ∞, respectively, except for p_2_, which was constrained to the range from 1 to ∞. All other parameters of the method were kept at their default values.

To estimate the error in estimating the percolation threshold, we performed 100 bootstrap iterations by randomly resampling 2,000 GEMs and fitting the linear-power law function for each resampled dataset. The error for the percolation threshold was reported as a standard deviation across the bootstrap iterations.

### Clustering of Metabolites and Pathway Analysis

For each carbon compound considered in the analysis, we calculated the probability of being in the largest component as a function of the number of network reactions. This profile was determined by calculating the moving average (using a window size of 300) of a binary variable describing the presence or absence of each carbon compound in the largest component across all GEMs, sorted by increasing network sizes. Based on the metabolites’ probability profiles, we performed k-means clustering using 5 clusters using the “kmeans” function in MATLAB. For visualization of the two most significant clusters of carbon compounds, we utilized the metabolism map provided by the interactive Pathways Explorer (iPath 3) tool (Darzi et al., 2018).

To identify the metabolic pathways represented in each cluster of carbon compounds, we mapped their KEGG IDs and conducted a Pathway Analysis using the MetaboAnalyst 6.0 web-based platform (Pang et al., 2024). We applied the hypergeometric test as the enrichment method, using the relative-betweenness centrality as the topology measure, and selected the pathway library of *Escherichia coli* K-12 MG1655 (KEGG) as the reference metabolome.

### Analysis of Bacterial Gene Content

The 2021 update of the COG database was used (Galperin et al., 2021). We categorized COGs into several functional groups. Specifically, housekeeping genes, including COG categories J, A, L, B, D, O, and K; and environmental genes, including COG categories V, X, and U. For each of the 1,310 microbial genomes in the COG database, we calculated the number of genes assigned to these COGs. Total counts of coding sequences (CDS) for each genome were obtained from NCBI annotations (Tatusova et al., 2016). We analyzed the number of transcription factors within each bacterial genome obtained from the KBase platform. To predict genes likely encoding transcription factors, we conducted Hidden Markov Models (HMMs) searches based on the PFAM database (Mistry et al., 2021) against the corresponding bacterial proteomes using the “pfam_scan.pl” tool. The options “translate” and “clan_overlap” were applied to generate the proteome from the genome and to allow overlapping hits within clan member families, respectively. Hits were identified based on an E-value ≤ 10^−3^ for members of a predefined list of 123 PFAM families previously described as likely DNA-binding transcription factors (Martinez-Liu et al., 2021)

For each functional category, we analyzed the relationship between the number of genes assigned to the category and the total number of CDS in the corresponding genome across bacteria. A piecewise linear regression was then fitted on the logarithmic scale; the breakpoint transition point for the fits was determined as the number of CDS in the genome approximately corresponding to the number of reactions at the percolation threshold (**Suppl Figure 2a**). The statistical significance of the piecewise versus single linear regression was assessed using an F-test, which compares the sum of squared errors of the piecewise model with that of the single linear model (Ludden et al., 1994).

### Analysis of Metagenomic Assemblies

Data on estimated genome sizes from metagenomic samples for species in terrestrial and aquatic environments were obtained from a published study (Rodríguez-Gijón et al., 2022). To assess the presence of bimodality in the size distributions, gamma mixture models with two components were fitted to the genome sizes distribution for each environment using an expectation-maximization algorithm (Dempster et al., 1977). The quality of the mixture models was evaluated using the Akaike Information Criterion (AIC) (Akaike, 1974).

## References

Akaike H. 1974. A new look at the statistical model identification. IEEE Trans Automat Contr 19:716–723.

Albert R, Jeong H, Barabási A-L. 2000. Error and attack tolerance of complex networks. Nature 406:378– 382.

Arkin AP, Cottingham RW, Henry CS, Harris NL, Stevens RL, Maslov S, Dehal P, Ware D, Perez F, Canon S, others. 2018. KBase: the United States department of energy systems biology knowledgebase. Nat Biotechnol 36:566–569.

Barabási A-L, Albert R. 1999. Emergence of scaling in random networks. Science 286:509–512.

Bentkowski P, Van Oosterhout C, Mock T. 2015. A model of genome size evolution for prokaryotes in stable and fluctuating environments. Genome Biol Evol 7:2344–2351.

Berg IA. 2011. Ecological aspects of the distribution of different autotrophic CO2 fixation pathways. Appl Environ Microbiol 77:1925–1936.

Burke GR, Moran NA. 2011. Massive genomic decay in Serratia symbiotica, a recently evolved symbiont of aphids. Genome Biol Evol 3:195–208.

Burstein D, Sun CL, Brown CT, Sharon I, Anantharaman K, Probst AJ, Thomas BC, Banfield JF. 2016. Major bacterial lineages are essentially devoid of CRISPR-Cas viral defence systems. Nat Commun 7:1–8.

Cottrell MT, Kirchman DL. 2016. Transcriptional control in marine copiotrophic and oligotrophic bacteria with streamlined genomes. Appl Environ Microbiol 82:6010–6018.

Csete M, Doyle J. 2004. Bow ties, metabolism and disease. Trends Biotechnol 22:446–450.

Darzi Y, Letunic I, Bork P, Yamada T. 2018. iPath3. 0: interactive pathways explorer v3. Nucleic Acids Res 46:W510–W513.

Dempster AP, Laird NM, Rubin DB. 1977. Maximum likelihood from incomplete data via the EM algorithm. Journal of the royal statistical society: series B (methodological) 39:1–22.

D’Souza G, Kost C. 2016. Experimental evolution of metabolic dependency in bacteria. PLoS Genet 12:e1006364.

Fisher RM, Henry LM, Cornwallis CK, Kiers ET, West SA. 2017. The evolution of host-symbiont dependence. Nat Commun 8:1–8.

Galperin MY, Wolf YI, Makarova KS, Vera Alvarez R, Landsman D, Koonin E V. 2021. COG database update: focus on microbial diversity, model organisms, and widespread pathogens. Nucleic Acids Res 49:D274–D281.

Gao Y, Yuan Q, Mao Z, Liu H, Ma H. 2021. Global connectivity in genome-scale metabolic networks revealed by comprehensive FBA-based pathway analysis. BMC Microbiol 21:1–15.

González-Domenech CM, Belda E, Patiño-Navarrete R, Moya A, Peretó J, Latorre A. 2012. Metabolic stasis in an ancient symbiosis: genome-scale metabolic networks from two Blattabacterium cuenoti strains, primary endosymbionts of cockroaches. BMC Microbiol 12:1–11.

Heirendt L, Arreckx S, Pfau T, Mendoza SN, Richelle A, Heinken A, Haraldsdóttir HS, Wachowiak J, Keating SM, Vlasov V, others. 2019. Creation and analysis of biochemical constraint-based models using the COBRA Toolbox v. 3.0. Nat Protoc 14:639–702.

Henry CS, DeJongh M, Best AA, Frybarger PM, Linsay B, Stevens RL. 2010. High-throughput generation, optimization and analysis of genome-scale metabolic models. Nat Biotechnol 28:977–982.

Herlemann DPR, Geissinger O, Ikeda-Ohtsubo W, Kunin V, Sun H, Lapidus A, Hugenholtz P, Brune A. 2009. Genomic analysis of “Elusimicrobium minutum,” the first cultivated representative of the phylum “Elusimicrobia”(formerly termite group 1). Appl Environ Microbiol 75:2841–2849.

Hugenholtz P, Tyson GW. 2008. Metagenomics. Nature 455:481–483. doi:10.1038/455481a

Jeong H, Tombor B, Albert R, Oltvai ZN, Barabási A-L. 2000. The large-scale organization of metabolic networks. Nature 407:651–654.

Keller MA, Kampjut D, Harrison SA, Ralser M. 2017. Sulfate radicals enable a non-enzymatic Krebs cycle precursor. Nat Ecol Evol 1:83.

Koonin E V, Wolf YI. 2008. Genomics of bacteria and archaea: the emerging dynamic view of the prokaryotic world. Nucleic Acids Res 36:6688–6719.

Land M, Hauser L, Jun S-R, Nookaew I, Leuze MR, Ahn T-H, Karpinets T, Lund O, Kora G, Wassenaar T, others. 2015. Insights from 20 years of bacterial genome sequencing. Funct Integr Genomics 15:141–161.

Lewis NE, Nagarajan H, Palsson BO. 2012. Constraining the metabolic genotype–phenotype relationship using a phylogeny of in silico methods. Nat Rev Microbiol 10:291–305.

Li M, Liu R-R, Lü L, Hu M-B, Xu S, Zhang Y-C. 2021. Percolation on complex networks: Theory and application. Phys Rep 907:1–68.

Lin Y-T, Ip JC-H, He X, Gao Z-M, Perez M, Xu T, Sun J, Qian P-Y, Qiu J-W. 2024. Scallop-bacteria symbiosis from the deep sea reveals strong genomic coupling in the absence of cellular integration. ISME J 18:wrae048.

Ludden TM, Beal SL, Sheiner LB. 1994. Comparison of the Akaike Information Criterion, the Schwarz criterion and the F test as guides to model selection. J Pharmacokinet Biopharm 22:431–445.

Madigan MT, Martinko JM, Stahl DA, Clark DP. 2012. Brock biology of microorganisms, Global Edition. San Francisco, TX: Pearson Benjamin Cummings.

Martinez-Liu L, Hernandez-Guerrero R, Rivera-Gomez N, Martinez-Nuñez MA, Escobar-Turriza P, Peeters E, Perez-Rueda E. 2021. Comparative genomics of DNA-binding transcription factors in archaeal and bacterial organisms. PLoS One 16:e0254025.

McCutcheon JP, Moran NA. 2012. Extreme genome reduction in symbiotic bacteria. Nat Rev Microbiol 10:13–26.

Mistry J, Chuguransky S, Williams L, Qureshi M, Salazar GA, Sonnhammer ELL, Tosatto SCE, Paladin L, Raj S, Richardson LJ, others. 2021. Pfam: The protein families database in 2021. Nucleic Acids Res 49:D412–D419.

Molina N, van Nimwegen E. 2009. Scaling laws in functional genome content across prokaryotic clades and lifestyles. Trends in genetics 25:243–247.

Murray GGR, Charlesworth J, Miller EL, Casey MJ, Lloyd CT, Gottschalk M, Tucker AW, Welch JJ, Weinert LA. 2021. Genome reduction is associated with bacterial pathogenicity across different scales of temporal and ecological divergence. Mol Biol Evol 38:1570–1579.

Nakabachi A, Yamashita A, Toh H, Ishikawa H, Dunbar HE, Moran NA, Hattori M. 2006. The 160-kilobase genome of the bacterial endosymbiont Carsonella. Science 314:267.

Nerima B, Nilsson D, Mäser P. 2010. Comparative genomics of metabolic networks of free-living and parasitic eukaryotes. BMC Genomics 11:1–8.

Nunoura T, Chikaraishi Y, Izaki R, Suwa T, Sato T, Harada T, Mori K, Kato Y, Miyazaki M, Shimamura S, others. 2018. A primordial and reversible TCA cycle in a facultatively chemolithoautotrophic thermophile. Science 359:559–563.

Oberhardt MA, Zarecki R, Gronow S, Lang E, Klenk H-P, Gophna U, Ruppin E. 2015. Harnessing the landscape of microbial culture media to predict new organism–media pairings. Nat Commun 6:1– 14.

Orth JD, Thiele I, Palsson BØ. 2010. What is flux balance analysis? Nat Biotechnol 28:245–248.

Pang Z, Lu Y, Zhou G, Hui F, Xu L, Viau C, Spigelman AF, MacDonald PE, Wishart DS, Li S, others. 2024. MetaboAnalyst 6.0: towards a unified platform for metabolomics data processing, analysis and interpretation. Nucleic Acids Res gkae253.

Parter M, Kashtan N, Alon U. 2007. Environmental variability and modularity of bacterial metabolic networks. BMC Evol Biol 7:1–8.

Passi A, Tibocha-Bonilla JD, Kumar M, Tec-Campos D, Zengler K, Zuniga C. 2021. Genome-scale metabolic modeling enables in-depth understanding of big data. Metabolites 12:14.

Plata G, Henry CS, Vitkup D. 2015. Long-term phenotypic evolution of bacteria. Nature 517:369–372.

Power JJ, Pinheiro F, Pompei S, Kovacova V, Yüksel M, Rathmann I, Förster M, Lässig M, Maier B. 2021. Adaptive evolution of hybrid bacteria by horizontal gene transfer. Proceedings of the national Academy of Sciences 118:e2007873118.

Pudlo NA, Urs K, Crawford R, Pirani A, Atherly T, Jimenez R, Terrapon N, Henrissat B, Peterson D, Ziemer C, others. 2022. Phenotypic and genomic diversification in complex carbohydrate-degrading human gut bacteria. mSystems 7:e00947–21.

Ravasz E, Somera AL, Mongru DA, Oltvai ZN, Barabási A-L. 2002. Hierarchical organization of modularity in metabolic networks. Science 297:1551–1555.

Rodríguez-Gijón A, Nuy JK, Mehrshad M, Buck M, Schulz F, Woyke T, Garcia SL. 2022. A Genomic Perspective Across Earth’s Microbiomes Reveals That Genome Size in Archaea and Bacteria Is Linked to Ecosystem Type and Trophic Strategy. Front Microbiol 12. doi:10.3389/fmicb.2021.761869

Sela I, Wolf YI, Koonin E V. 2019. Selection and Genome Plasticity as the Key Factors in the Evolution of Bacteria. Phys Rev X 9:31018. doi:10.1103/PhysRevX.9.031018

Serrano MÁ, Boguná M, Sagués F. 2012. Uncovering the hidden geometry behind metabolic networks. Mol Biosyst 8:843–850.

Smart AG, Amaral LAN, Ottino JM. 2008. Cascading failure and robustness in metabolic networks. Proceedings of the National Academy of Sciences 105:13223–13228.

Stauffer D. 1979. Scaling theory of percolation clusters. Phys Rep 54:1–74.

Stobbe MD, Houten SM, van Kampen AHC, Wanders RJA, Moerland PD. 2012. Improving the description of metabolic networks: the TCA cycle as example. The FASEB Journal 26:3625–3636.

Tatusova T, DiCuccio M, Badretdin A, Chetvernin V, Nawrocki EP, Zaslavsky L, Lomsadze A, Pruitt KD, Borodovsky M, Ostell J. 2016. NCBI prokaryotic genome annotation pipeline. Nucleic Acids Res 44:6614–6624.

Toh H, Weiss BL, Perkin SAH, Yamashita A, Oshima K, Hattori M, Aksoy S. 2006. Massive genome erosion and functional adaptations provide insights into the symbiotic lifestyle of Sodalis glossinidius in the tsetse host. Genome Res 16:149–156.

Wolf YI, Koonin E V. 2013. Genome reduction as the dominant mode of evolution. Bioessays 35:829– 837.

Yang S. 2013. Networks: An Introduction by MEJ Newman: Oxford, UK: Oxford University Press. 720 pp., 85.00.

Zarecki R, Oberhardt MA, Reshef L, Gophna U, Ruppin E. 2014. A novel nutritional predictor links microbial fastidiousness with lowered ubiquity, growth rate, and cooperativeness. PLoS Comput Biol 10:e1003726.

Zeng X, Xing X, Gupta M, Keber FC, Lopez JG, Lee Y-CJ, Roichman A, Wang L, Neinast MD, Donia MS, others. 2022. Gut bacterial nutrient preferences quantified in vivo. Cell 185:3441–3456.

